# The androgen receptor does not directly regulate the transcription of DNA damage response genes

**DOI:** 10.1101/2023.05.13.540653

**Authors:** Sylwia Hasterok, Thomas G. Scott, Devin G. Roller, Adam Spencer, Arun B. Dutta, Kizhakke M Sathyan, Daniel E. Frigo, Michael J. Guertin, Daniel Gioeli

**Author notes:** ^#^ These authors contributed equally to the work (Sylwia Hasterok, Thomas G. Scott). To whom correspondence should be addressed. Daniel Gioeli, Department of Microbiology, Immunology, and Cancer Biology, University of Virginia, Charlottesville, Virginia, 22908, United States of America.

## Abstract

The clinical success of combined androgen deprivation therapy (ADT) and radiation therapy (RT) in prostate cancer (PCa) created interest in understanding the mechanistic links between androgen receptor (AR) signaling and the DNA damage response (DDR). Convergent data have led to a model where AR both regulates, and is regulated by, the DDR. Integral to this model is that the AR regulates the transcription of DDR genes both at steady state and in response to ionizing radiation (IR). In this study, we sought to determine which immediate transcriptional changes are induced by IR in an AR-dependent manner. Using PRO-seq to quantify changes in nascent RNA transcription in response to IR, the AR antagonist enzalutamide, or the combination of the two, we find that enzalutamide treatment significantly decreased expression of canonical AR target genes but had no effect on DDR gene sets in PCa cells. Surprisingly, we also found that the AR is not a primary regulator of DDR genes either in response to IR or at steady state in asynchronously growing PCa cells. Our data indicate that the clinical benefit of ADT and RT is not due to the direct regulation of DDR gene transcription by AR.

## Introduction

Prostate cancer (PCa) continues to rank as one of the leading cancers in men worldwide (1). PCa growth and survival are highly dependent on androgen receptor (AR) signaling (2). AR activation is a major driving force behind not only the development and progression of androgen-dependent PCa but also the progression to incurable, lethal, castration-resistant prostate cancer (CRPC) (3). To date, standard of care treatments for advancing PCa range from surgery, chemotherapy, radiation therapy (RT), androgen deprivation therapy (ADT), and combinations thereof (4).

Since there is a demonstrated clinical benefit for combining ADT with RT for PCa, many studies have explored the link between the DNA damage response (DDR) and AR (5,6). There are multiple connections between the AR and the DDR, culminating in a model where the AR both regulates, and is regulated by, the DDR. Key studies a decade ago suggested the model that activation of the AR by ionizing radiation (IR) increases the transcription of DDR genes, leading to rapid repair of DNA damage and promoting radioresistance in PCa cells (7–9). These papers provided evidence that radiation-induced DNA damage enhances the AR-mediated expression of genes involved in DNA damage repair. Additionally, androgens regulate the expression of the catalytic subunit of DNA-PK, a critical component of non-homologous end joining that is responsible for engaging repair factors at sites of DNA damage, promoting the resolution of double-stranded breaks and leading to resistance to IR genotoxicity. These findings were supported by subsequent studies indicating androgen and AR regulation of subsets of DDR genes (10,11). In one study, the specific DNA repair genes regulated by AR were particular to the model system and disease state examined (10).

Other studies have provided evidence that the AR is regulated by DDR components. PARP-1 was recruited to AR binding sites, enhancing AR occupancy and transcriptional function (12). Tandem mass spectroscopy analysis identified Ku70 and Ku80 as direct AR-interacting proteins that positively regulate AR transactivation (13). Furthermore, BRCA1 physically interacted with the DNA-binding domain (DBD) of AR to enhance AR transactivation and induce androgen-mediated cell death through p21 expression (14). In contrast, the association of the ligand binding domain (LBD) of AR with hRad9, a crucial member of the checkpoint Rad family, suppressed AR transactivation by preventing the androgen-induced interaction between the n-terminus and c-terminus of AR (15). Other groups reported non-genomic effects as a result of DDR protein-AR interactions. Mediator of DNA damage checkpoint protein 1 (MDC1), an essential player in the Intra-S phase and G2/M checkpoints, physically associated with full-length AR (FL-AR) and the AR splice variant v7 (AR^V7^) to negatively regulate PCa cell growth and migration (16). Another study showed that increased clonogenic survival following IR was a consequence of DNA-PKc directly complexing with both FL-AR and AR^V5-7^, with radiation increasing these interactions and enzalutamide blocking the association with FL-AR but not AR^V5-7^ (17). Collectively, these data provide evidence that the AR is integrated into the DDR, interfacing at multiple points, and support an integrated mechanistic model of cooperation between the AR and DDR machinery to explain the clinically observed cooperativity of ADT and RT.

Our own work previously demonstrated a connection between AR and CHK2, a cell cycle checkpoint regulator downstream of DNA damage and replication blockade (18,19). We found that IR is a major driver of the CHK2**–**AR interaction and that binding of CHK2 to the AR antagonizes its function leading to AR inactivation. Moreover, alterations in CHK2 activity, caused by gene mutations or loss of expression, are common in PCa and support increased AR activity. We proposed that this, in turn, stimulates the DDR machinery, which repairs IR-induced DNA lesions and promotes PCa cell survival. As part of our study, we tested the impact of the CHK2**–**AR interaction on the transcription of ten different DDR genes previously defined as part of an AR-associated DNA repair gene signature (8). To our surprise, we detected only a modest induction by IR or androgen treatment in just two of the ten DDR genes. This inconsistency aligned with the earlier report indicating that the AR-regulated DDR genes may depend on the model system, experimental conditions, and disease state examined (10). This suggests that the model of AR regulating DDR genes is incomplete and needs further examination. One limitation of the aforementioned studies is that transcripts were generally measured with delayed kinetics (typically one to four days) following treatment with hormone, anti-androgens, or IR. Thus, the androgen-responsive genes identified in these studies could include those that are downstream from primary AR targets. We therefore sought to determine the primary AR transcriptional targets induced by IR to uncover missing mechanistic details for how the AR regulates the DDR.

## Results

### AR does not regulate the primary IR-induced DNA damage transcriptional response

To determine if AR is activated by DNA damage, we measured IR-induced changes in nascent RNA transcription in asynchronously growing androgen-responsive LNCaP cells. We performed precision nuclear run-on sequencing (PRO-seq) on LNCaP cells grown in whole medium 1 hour after treatment with 6 Gray (Gy) of IR or a mock treatment (Figure 1A) to identify the primary transcriptional response to IR. This dose of IR was able to induce TMPRSS2, and to a lesser extent FKBP5, at 24hrs in LNCaP and C4-2 cells, recapitulating what was previously reported for C4-2 cells (Figure 1B) (20). We observed an immediate transcriptional response to the IR-induced DNA damage, with 218 genes significantly activated and 80 genes significantly repressed at a false discovery rate (FDR) of 0.1 (Figure 1C-D; Supplemental Figure 1A; Supplemental Table 1). Using gene set enrichment analysis (GSEA) with the Hallmark Gene Sets (21), we explored which biological processes were over-represented in the IR-induced DNA damage transcriptional response (Figure 1E). At an FDR of 0.05, genes in the P53 Pathway were predominately activated, and genes in the Mitotic Spindle and G2/M Checkpoint gene sets tended to be repressed in response to IR. The Hallmark Androgen Response gene set was not identified at an FDR of 0.05 or when the FDR was relaxed to 0.2, suggesting that AR is not directly activated by IR. The Hallmark Androgen Response gene set has an adjusted p-value of 0. Article Tracked Changes. Consistent with AR not being activated in response to IR, we do not observe significant expression changes of seven canonical AR target genes: KLK3, KLK2, TMPRSS2, NKX3.1, FKBP5, SGK1, and STEAP4 (Supplemental Table 2). To identify candidate sequence-specific transcription factors (TFs) that drive the IR-induced DNA damage transcriptional response, we first defined putative regulatory elements (e.g. enhancers and promoters) with dREG (22). We identified putative regulatory elements responsive to IR with DESeq2 (Supplemental Figure 1B). Next, we performed *de novo* motif analysis using MEME (23) within the increasing putative regulatory elements and matched these motifs to the JASPAR database (24), and found only the p53 motif (Figure 1F), consistent with the GSEA. We found no motifs within the decreasing putative regulatory elements. Using FIMO (25) to identify p53 motif instances genome-wide, we found a significant enrichment (Chi-squared test p-value < 2.2 x 10^-16^) of p53 motifs in IR-activated putative regulatory elements, whereas there was no enrichment of p53 motifs in the IR-repressed putative regulatory elements (Figure 1G). These data indicate that the predominant primary transcriptional response triggered by IR in LNCaP PCa cells is driven by p53, consistent with the canonical p53 DNA damage transcriptional response. When we explicitly identified androgen response element (ARE) motif instances genome-wide, we found a slight enrichment in both the IR-activated and the IR-repressed putative regulatory elements (Figure 1H). This suggests that if changes in gene expression induced by IR are mediated by the AR, there are a comparable number of AR-regulated genes that are repressed as are activated. Even though the majority of the transcriptional response to IR is activation of transcription, there does not appear to be a genome-wide increase in AR activity.

**Figure 1.**
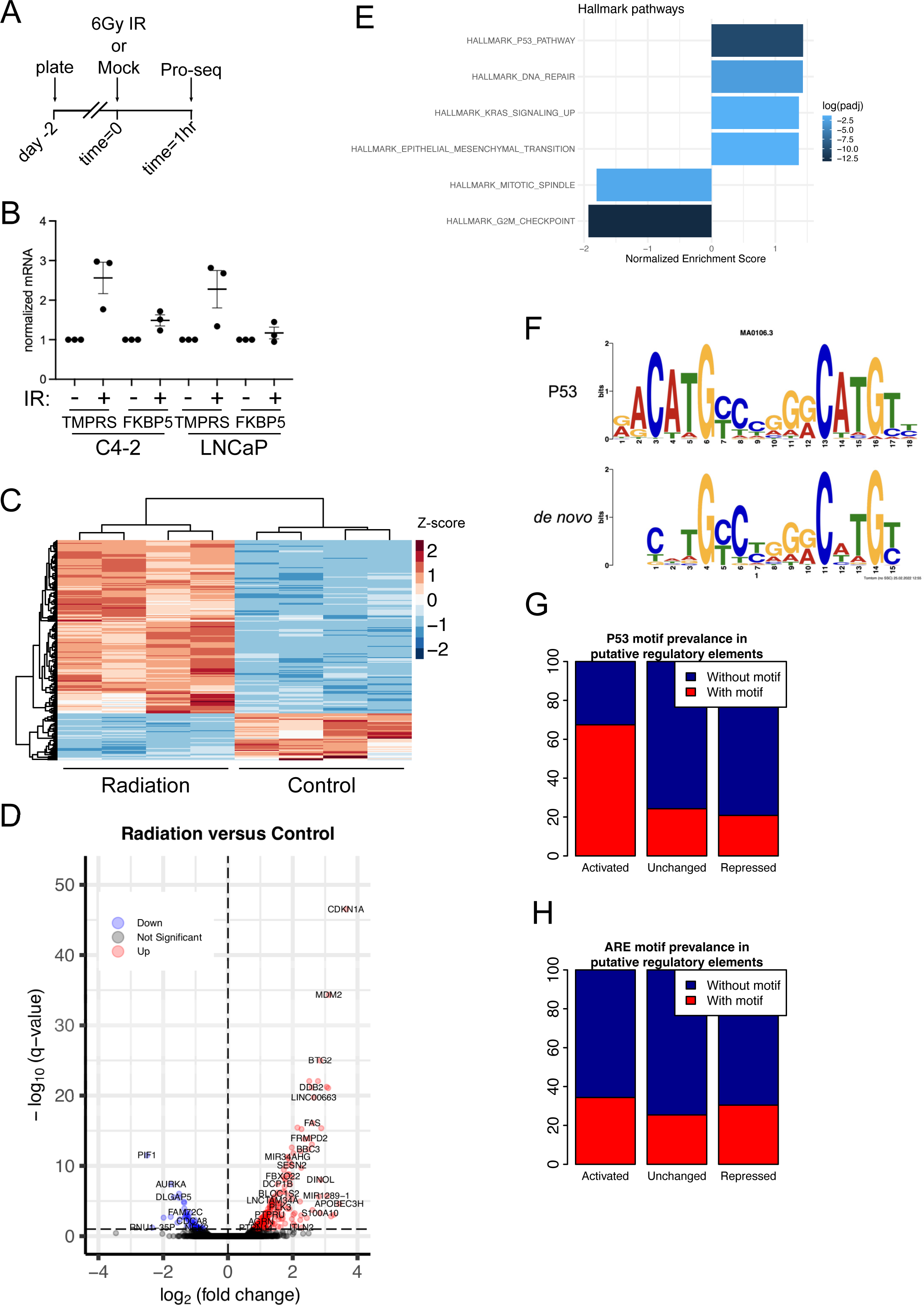
IR activates a p53 transcriptional response in PCa cells. (A) Outline of experiment; n=4 per condition. (B) Validation of IR induction of TMPRSS2 (TMPRS) and FKBP5 in C4-2 and LNCaP cells at 24hrs following 6Gy. (C) Heat map of PRO-seq signal reflecting gene expression values, scaled by row. (D) Volcano plot highlighting differentially expressed genes upon radiation treatment. The horizontal dashed black line denotes an adjusted p-value of 0.1. Points having a positive or negative fold-change are shown in red (up in response to radiation) or blue (down in response to radiation), respectively. (E) GSEA of IR-induced primary transcriptional response. All significant (FDR 0.05) sets are shown. (F) *de novo* identified motif from IR-ctivated putative regulatory elements matching the canonical p53 motif from the JASPAR database. (G) Differential p53 motif prevalence in putative regulatory elements that change in response to radiation. (H) Differential AR motif prevalence in putative regulatory elements that change in response to radiation.

To explicitly identify AR-regulated transcripts among the IR-induced DNA damage primary transcriptional response we pretreated LNCaP PCa cells with 10 µM enzalutamide for 30 minutes prior to 6 Gy IR and then performed PRO-seq following a 1-hour incubation post-IR (Figure 2A). The 10 µM dose of enzalutamide maximally inhibited LNCaP PCa cell growth (Figure 2B). An identical maximum inhibition was achieved with darolutamide, another second-generation antiandrogen that is structurally distinct from enzalutamide. For additional controls, we confirmed that AR antagonism with enzalutamide was radiosensitizing and AR activation with the synthetic androgen R1881 increased cell growth in the presence of IR (Figure 2C). Hierarchical clustering of the dynamic genes (Figure 2D) segregates treatment groups first by IR and then by enzalutamide, reinforcing that IR is the major driver of gene expression changes relative to enzalutamide. The volcano plots (Figure 2E) illustrate a robust transcriptional response to IR, comparable to what we observed earlier (Figure 1D). See Supplemental Figure 1 for corresponding MA plots and Supplemental Tables 3 and 4 for the list of gene expression changes in response to IR and enzalutamide, respectively. Enzalutamide treatment significantly decreased expression of 182 genes (FDR 0.05); GSEA reveals that these genes represent canonical AR target genes (Hallmark Androgen Response). We find no Hallmark or Reactome gene set signature of DNA damage within genes significantly changed by enzalutamide (at FDR 0.05 or 0.2).

**Figure 2.**
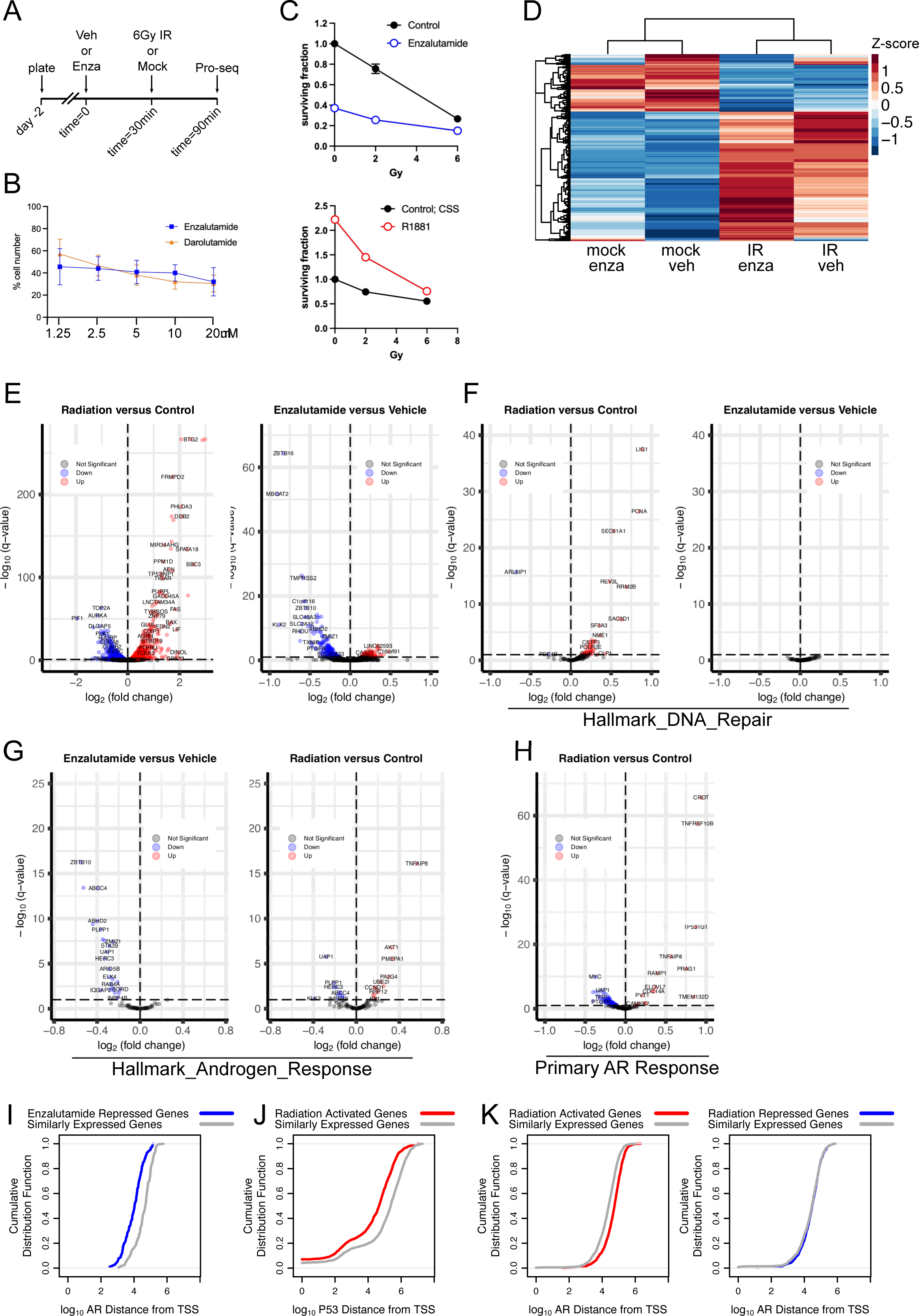
The directly AR-regulated transcriptome is not a major contributor to the immediate transcriptional response to DNA damage. (A) Outline of experiment; n=2 per condition. (B) Validation of growth inhibition by AR antagonism in LNCaP cells. Percent cells relative to vehicle control following 6 days. (C) Validation of radiosensitization with AR antagonism using 10µM enzalutamide and radioprotection with AR activation using 0.05nM R1881. (D) Heat map of PRO-seq signal reflecting gene expression values, scaled by row for control, radiation (6Gy), enzalutamide (10µM), and radiation+enzalutamide (6Gy+10µM enzalutamide) samples. (E) Volcano plot highlighting differentially expressed genes upon radiation treatment (left) or enzalutamide treatment (right). (F) Volcano plot highlighting differentially expressed genes in the Hallmark DNA Repair gene set upon radiation treatment (left) or enzalutamide treatment (right). (G) Volcano plot highlighting differentially expressed genes in our primary AR target gene set upon enzalutamide treatment (left) or radiation treatment (right). (H) Volcano plot highlighting differentially expressed genes in the Hallmark DNA Repair gene set upon radiation treatment. In (E-H), the horizontal dashed black line denotes an adjusted p-value of 0.1. Points having a positive or negative fold-change are shown in red (up in response to radiation or enzalutamide) or blue (down in response to radiation or enzalutamide), respectively. (I-K) Cumulative Distribution Function plots (Kolmogorov–Smirnov test) of the distances from the TSS of (I) IR-activated genes to AR binding sites, (J) enzalutamide-repressed genes to AR binding sites, and (K) IR-activated genes to p53 binding sites.

To further test if AR is part of the primary transcriptional response to IR-induced DNA damage, we explicitly examined the changes in expression of the genes in the Hallmark DNA Repair gene set upon either IR or enzalutamide (Figure 2F). IR significantly activated multiple genes, while enzalutamide treatment did not significantly change expression of any genes in the set. We also explicitly surveyed the PRO-seq data for changes in expression of the DDR genes previously defined as AR-regulated (7,8,10) (Supplemental Table 5). We did not find any significant changes in response to enzalutamide in the previously defined 32 direct DDR AR target genes (8), or in DNA-PK (7), and only four of 36 previously defined androgen-regulated DDR genes from LNCaP (10) were significantly changed by enzalutamide. Thus, in asynchronously growing PCa cells the AR does not appear to directly regulate the expression of genes in the DDR, even when cells are exposed to IR DNA damage. Next, we examined the effect of enzalutamide and IR on the expression of genes in the Hallmark Androgen Response gene set (Figure 2G). As expected, enzalutamide treatment had a significant effect on many genes in the set. IR both activated and repressed genes in the Hallmark Androgen Response set, indicating that IR does not broadly activate or inhibit AR at all AR target genes. To test this further, we defined a Primary AR Target gene set from our own PRO-seq data of enzalutamide treated LNCaP cells and examined the effect of IR on the expression of these genes (Figure 2H). As with the genes in the Hallmark Androgen Response gene set, expression of Primary AR Target genes both increase and decrease in response to IR, consistent with TFs other than AR regulating gene expression changes in response to IR. Finally, we assessed the proximity between the transcription start site (TSS) of each gene regulated by IR or enzalutamide to AR or p53 binding sites, as defined by publicly available Chromatin Immunoprecipitation (ChIP) sequencing peaks; AR-ChIP peaks were from LNCaP cells while the p53 peaks were from A549 cells (26,27). As expected, enzalutamide-repressed genes tend to be closer to AR binding sites, indicating that these genes are direct targets of AR activation at baseline (Figure 2I). Similarly, IR-activated genes tend to be closer to p53 binding sites, consistent with a mechanism in which p53 activates a substantial proportion of these genes in response to IR (Figure 2J). In contrast, IR-responsive genes are not significantly closer to AR binding sites than IR-unchanged genes are, suggesting that AR is not responsible for the IR-responsive transcriptional changes (Figure 2K) and that TFs other than AR drive changes in gene expression induced by IR. Collectively these data indicate that the AR is not part of a primary IR-induced DNA damage transcriptional response in LNCaP PCa cells, nor is AR a primary regulator of genes in the DDR under normal growth conditions. This is further corroborated by analysis of published GRO-seq data of hormone-stimulated LNCaP and VCaP cells (28,29); neither of these studies of the hormone-induced primary AR transcriptional response identified gene sets associated with the DDR. Furthermore, we previously tested for changes in AR S81 phosphorylation in response to IR and found no change in S81 phosphorylation (19). Since the predominant kinase that phosphorylates S81 is CDK9, part of P-TEFb, and S81 phosphorylation is associated with AR transcriptional activity (30–32), the lack of IR induction of AR S81 phosphorylation is consistent with IR not activating the AR (19).

### IR activates a small subset of primary AR target genes

We tested for any role that AR may play in the transcriptional response to IR-induced DNA damage by identifying groups of genes that are regulated by both enzalutamide and IR. We find 32 genes activated by both enzalutamide and IR (Supplemental Figure 2A), 66 genes repressed by both enzalutamide and IR (Supplemental Figure 2B), 7 genes activated by enzalutamide and repressed by IR (Supplemental Figure 2C), and 16 genes activated by IR and repressed by enzalutamide (Figure 3A). This last group of 16 genes fits the pattern predicted for genes activated by AR in response to DNA damage. Over-representation analysis (ORA) for Hallmark gene sets indicated that the set of 32 genes activated by both enzalutamide and IR is enriched for cholesterol homeostasis genes, and the set of 66 genes repressed by both enzalutamide and IR are enriched for genes involved in androgen response and for genes downregulated by UV radiation (Supplemental Figure 2D); no gene sets were identified for the 7 genes activated by enzalutamide and repressed by IR. ORA indicated that the set of 16 genes activated by IR and repressed by enzalutamide is enriched for the canonical Hallmark Androgen Response gene set (Supplemental Figure 2D) but not for any gene sets implicated in the DDR. Further examining this set of genes, we found that alterations of these 16 genes in the TCGA PanCancer Atlas were associated with a poorer progression-free survival (PFS) (Supplemental Figure 2E). Interestingly, two of these 16 genes, TLE1 and WNT7B, were individually associated with a poorer PFS. This suggests that although there is only a small subset of primary AR target genes activated by IR, these genes may contribute to PCa progression.

**Figure 3.**
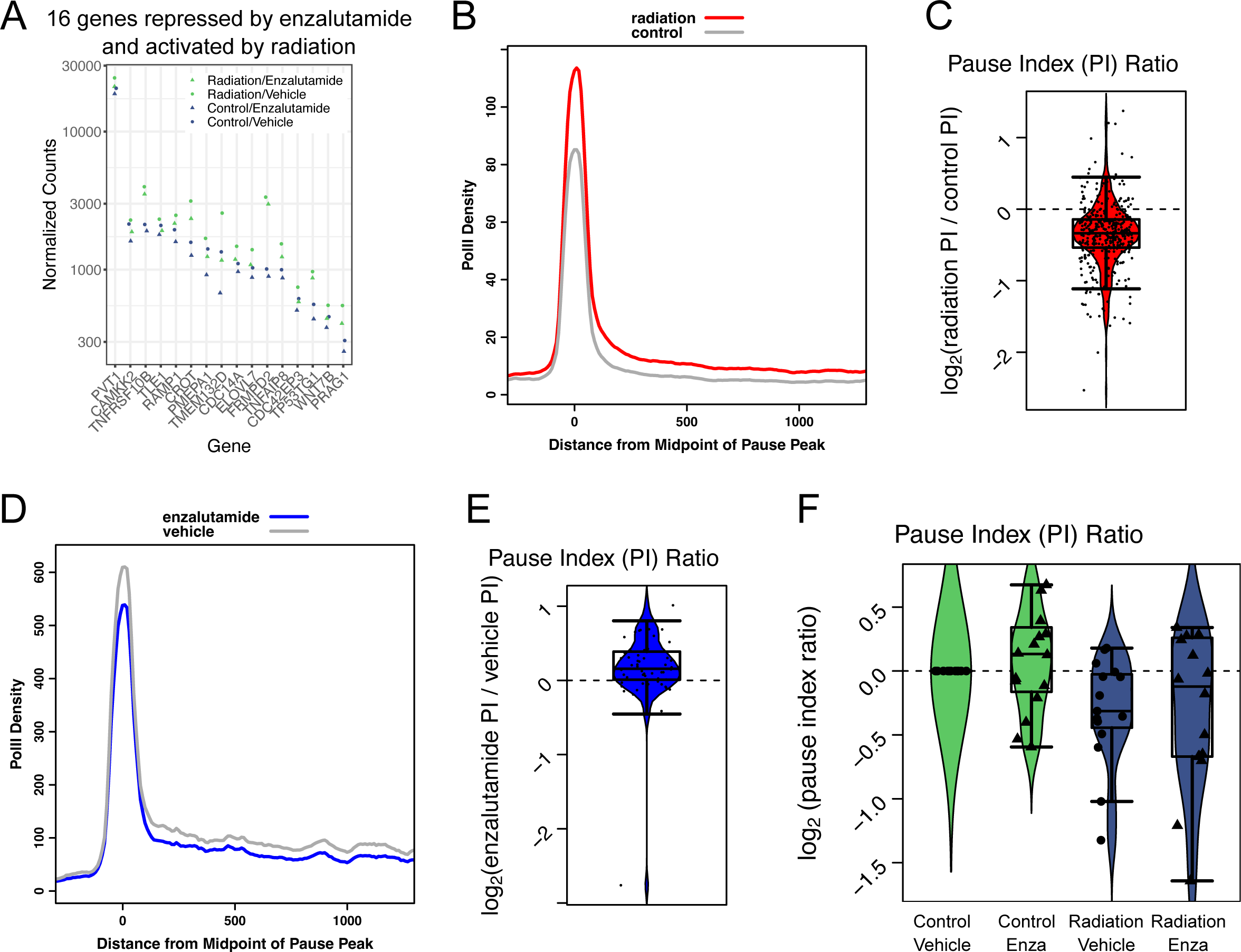
Enzalutamide treatment prevents Pol II promoter-proximal pause release triggered by IR within a subset of genes. (A) Normalized count expression data for genes activated by radiation and repressed by enzalutamide. (B) Pol II density plot of radiation-activated genes in radiation and control conditions. (C) Box and whisker and violin plot of the log-transformed ratio of pause index (PI) for radiation-activated genes in radiation and control conditions. (D) Pol II density plot of enzalutamide-repressed genes in enzalutamide and vehicle conditions. (E) Box and whisker and violin plot of the log-transformed ratio of PI for enzalutamide-repressed genes in enzalutamide and vehicle conditions. (F) Box and whisker and violin plot of the log-transformed ratio of PI for the 16 genes in (A) for the different treatments relative to the control and vehicle condition.

### IR increases, and enzalutamide decreases, promoter-proximal Pol II pause release

We next examined RNA polymerase II (Pol II) localization within genes responsive to IR, enzalutamide, or both treatments to determine if IR and AR regulate the same step of the Pol II transcription cycle (e.g. initiation, promoter-proximal pausing, elongation, or termination) (33,34). PRO-seq measures the genome-wide location of transcriptionally engaged Pol II at single-nucleotide resolution (35). This spatial precision allows us to determine Pol II density within the promoter-proximal pause region as well as within the gene body and to calculate a Pause Index (PI) for each gene using the ratio of these two densities (36). IR treatment leads to an increase in Pol II density in both regions within IR-activated genes (Figure 3B). Calculating the ratio of PI for each of these genes in IR relative to control conditions, we find that IR decreases PI, indicating that IR increases the rate of Pol II pause release (Figure 3C). We find that enzalutamide treatment leads to a decrease in Pol II density in both regions within enzalutamide-repressed genes and an increase in PI (Figures 3D and 3E), indicating that enzalutamide decreases the rate of Pol II pause release. These data would suggest that enzalutamide treatment may prevent the release of promoter-proximally paused Pol II on the 16 genes activated by IR and repressed by enzalutamide. Analysis of the changes in PI for the 16 genes is consistent with this prediction (Figure 3F); enzalutamide treatment increased PI, IR decreased PI, and the combined treatment led to an intermediate change in PI approximating zero. However, using only 16 genes is underpowered for statistical analysis of PI changes between conditions. Overall, these data indicate that, for the subset of primary AR target genes activated by IR, enzalutamide treatment may prevent the release of promoter-proximally paused Pol II in response to IR.

### Cellular controls mitigate the downstream effects of the IR-induced AR primary target gene subset

To explore these IR-activated enzalutamide-repressed genes further, we ranked the genes by the magnitude of enzalutamide inhibition of the IR induction below control levels and selected the top half to examine expression levels between normal and tumor tissue. This subset of eight genes was also associated with a poorer PFS in the TCGA PanCancer Atlas (Supplemental Figure 2F). We used Gene Expression Profiling Interaction Analysis (GEPIA2) analyzing the RNA sequencing expression data from the TCGA and the GTEx projects (37). We find that CAMKK2, ELOVL7, and PMEPA1 expression increases in tumor compared to control (Figure 4A). We dropped TMEM132D from further analysis since mRNA expression is very low in the prostate and PCa, and the TMEM132D protein does not seem to be expressed (38). We reasoned that, if AR activation was important in the cell response to IR, then the mRNA levels of these genes would remain elevated over time. Moreover, in previous studies on AR regulation of DDR genes (8,20), the earliest timepoint reported that showed a change in mRNA of a DDR gene was 24hrs, a result we confirmed in our cells (Figure 1B). Thus, despite the cell cycle arrest of PCa cells in response to IR or androgen deprivation (20) there is precedent for observing increased mRNA at 24hrs following IR. We assessed at 24 hours in LNCaP (Figure 4B), C4-2, 22Rv1, and VCaP cells (Supplemental Figure 3) the mRNA levels of CAMKK2, CDC42EP3, ELOVL7, PMEPA1, PVT1, TLE1, and WNT7B in response to 10 μM enzalutamide, 6 Gy IR, or a 30-minute pretreatment with 10 μM enzalutamide followed by 6 Gy IR. LNCaP and C4-2 cells are p53 wild-type, whereas 22Rv1 and VCaP are mutant p53. In LNCaP cells at 24 hours, enzalutamide significantly reduced the expression of five of the seven genes examined: CAMKK2, CDC42EP3, ELOVL7, PMEPA1, and WNT7B. IR significantly induced the expression of ELOVL7, PVT1, and TLE1. The mRNA abundance in the combination of enzalutamide and IR was significantly less than in IR alone for CAMKK2, ELOVL7, and WNT7B. Only ELOVL7 showed the same expression pattern at 1 and 24 hours post IR of a significant induction of expression by IR that is significantly antagonized by enzalutamide (Figures 3A and 4B). The expression patterns in C4-2, 22Rv1, and VCaP cells were appreciably different than in LNCaP cells, with only ELOVL7 and PVT1 in C4-2 and TLE1 in 22Rv1 cells increasing with IR. This could be explained in part due to the differences in p53 status. In C4-2 for CAMKK2 and ELOVL7, and in 22Rv1 for TLE1, the IR induction was antagonized by enzalutamide, consistent with these genes being induced by IR in an AR-dependent manner. The difference in response of these genes to IR and AR antagonism in the four PCa cell lines suggests that cell line heterogeneity affects the determination of any set of genes that increases with IR in an AR-dependent manner. Since ELOVL7 was the only gene with a significant induction of mRNA expression by IR that was antagonized by enzalutamide at both early (1 hour) and late (24 hour) timepoints in LNCaP and a second PCa cell line (C4-2; Supplemental Figure 3C), we next tested for changes in ELOVL7 protein expression over time in response to the same treatments (Figure 4C). We found no significant changes in protein expression from 15 minutes to 6 hours in response to IR, enzalutamide, or the combination thereof (Figure 4D). This suggests that, although the AR may activate transcription of a small set of genes in direct response to IR-induced DNA damage, additional cellular controls mitigate downstream effects of that activation, preventing sustained changes in mRNA or protein levels.

**Figure 4.**
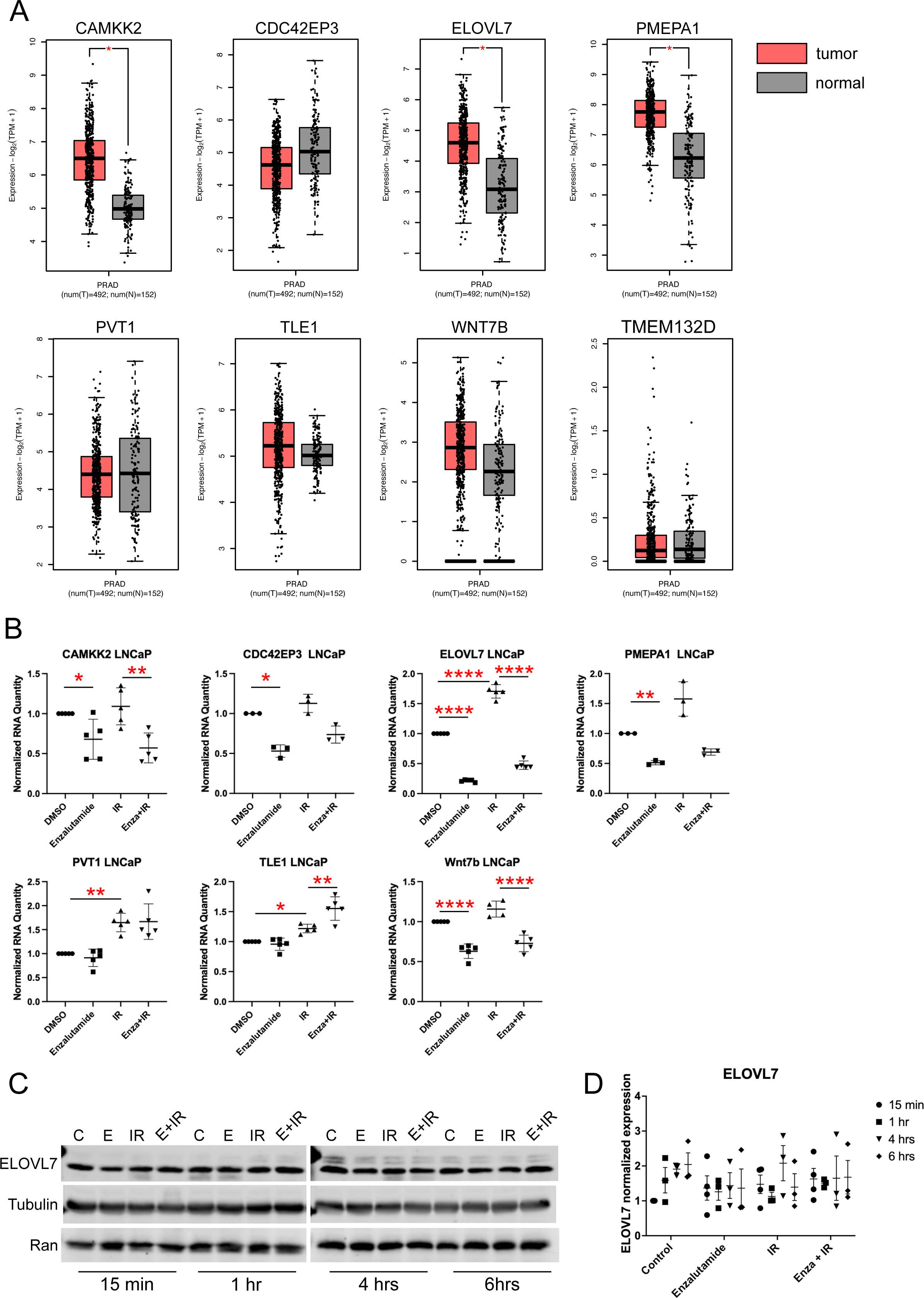
Regulation of candidate AR-regulated DDR genes in PCa tumors and LNCaP cells. (A) GEPIA2 analysis of the PCa TCGA and GTEx data of the top eight genes using a Log2FC cut off of 1 and a p-value cutoff of 0.01 (* <0.01). (B) Normalized mRNA expression of CAMKK2, CDC42EP3, ELOVL7, PMEPA1, PVT1, TLE1, and WNT7B at 24hrs in response to 6Gy radiation, 10µM enzalutamide, or combination thereof. * <0.05, ** <0.005, **** <0.0001 indicating a statistically significant effect of enzalutamide on control or IR, using ANOVA followed by Sidak’s multiple comparisons test. (C) Western blot data of ELOVL7, and loading controls Tubulin and Ran, over time (15 minutes, 1 hour, 4 hours, and 6 hours) in response to 6Gy radiation (IR), 10µM enzalutamide (E), or combination thereof (E+IR), and control (C) vehicle treated cells. (D) Quantitation of ELOVL7 expression normalized to Tubulin and Ran, with control at 15 minutes set to zero, n= 3 to 4.

### ELOVL7 and WNT7B knockdown leads to cell death while CAMKK2 knockout does not affect PCa cell growth under normal growth conditions regardless of IR or AR activity

Since alterations in these genes are associated with a shorter time of PFS (Supplemental Figure 2F), we next tested the impact of altering expression of these genes on PCa cell growth (Supplemental Figure 4). We selected CAMKK2, ELOVL7, and WNT7B as representatives of this gene set; all three have been implicated in PCa (39–42) and ELOVL7 is the only gene from the subset induced by IR and antagonized by enzalutamide at 1 and 24 hrs. Knockdown of ELOVL7 had a profound effect on cell growth, leading to clearing of cells from the knockdown wells (data not shown). Given the cell death induced by ELOVL7 knockdown it was not a surprise to see that enzalutamide treatment had no additional effect, and hormone treatment did not rescue growth when ELOVL7 was knocked down. The effect of WNT7B knockdown was similar to that of ELOVL7 but the effect was not as great. C4-2 cells had the same CAMKK2 expression pattern in response to enzalutamide and IR as LNCaP. Consistent with CAMKK2’s role in metabolic fitness (43), under nutrient-replete conditions, CAMKK2 knockout in C4-2 cells had no effect on *in vitro* growth, or response to enzalutamide or R1881 in our assay. These findings are in contrast to the established role of CAMKK2 in CRPC *in vivo* tumor growth (39), and further indicate that additional stresses are needed to reveal CAMKK2’s pro-cancer roles.

We next tested the impact of reduced expression of CAMKK2, ELOVL7, and WNT7B on cell response to IR in the presence and absence of hormone and enzalutamide (Supplemental Figure 5). For ELOVL7, knockdown reduced cell survival to the point that IR had no additional effect on survival and androgen offered no protection from the knockdown of ELOVL7. Knockdown of WNT7B also reduced cell survival, and IR had little additional effect. These results are consistent with ELOVL7 and WNT7B being essential for cell growth, which may account for why ELOVL7 protein levels do not change in response to IR and enzalutamide, despite changes in mRNA. In C4-2 cells the protective effect of AR activation is minimal, and while CAMKK2 knockout reduced survival overall, there was no sensitization to IR, and R1881 offered minimal protection. We also tested if knockout or knockdown of CAMKK2, ELOVL7, and WNT7B cooperated with enzalutamide and/or IR to reduce cell viability. Parallel effects with AR activation were observed for all three: ELOVL7 knockdown inhibited survival completely preventing any evaluation of cooperativity with enzalutamide; WNT7B knockdown further reduced survival in response to enzalutamide and at low dose IR; and CAMKK2 knockout did not generate enzalutamide sensitivity in C4-2 cells. These data support the conclusion that ELOVL7 and WNT7B expression are generally important for PCa cell growth and that revealing the role of CAMKK2 in PCa growth requires growth factor stress (39).

### ZBTB10 overexpression is not sufficient to rescue the radiosensitization of AR inhibition

A distinguishing feature of our data in comparison to prior work that has indicated that AR regulates the transcription of DDR genes (7,8,10,11) is that, by measuring nascent transcription shortly after enzalutamide treatment, we isolated the primary effects of AR inhibition. Downstream from these primary effects, AR may indirectly regulate DDR genes by activating the transcription of additional TFs that directly regulate DDR genes. Enzalutamide treatment significantly (adjusted p-value <0.05) repressed 17 TFs (Figure 5A). The most enzalutamide-repressed TF in the Hallmark Androgen Response is ZBTB10. In the TCGA PCa data, ZBTB10 amplification is associated with a lower disease-free survival (Figure 5B), and ZBTB10 expression is higher (p-value < 0.01) in tumor than normal (Figure 5C). In LNCaP and C4-2 cells at 24 hours, there was a significant induction of ZBTB10 expression by IR that was significantly antagonized by enzalutamide (Figure 5D, E) suggesting that ZBTB10 activation by AR may be important in the cellular response to IR. There was no effect of IR or enzalutamide on ZBTB10 expression in 22Rv1 cells. In VCaP cells, enzalutamide inhibited ZBTB10 expression, but expression was not affected by IR. We reasoned that, if AR activation of ZBTB10 expression mediated downstream activation of the previously reported DDR genes trigged by AR response to IR, then overexpression of ZBTB10 should lead to an increase in DDR gene expression, reduce the radiosensitization of enzalutamide, and enhance cell growth, even in the presence of IR. However, ZBTB10 overexpression (Figure 5F, G) did not lead to an increase in PRKDC, XRCC2, or XRCC3 (Figure 5H), or diminish the radiosensitization of enzalutamide (Figure 5I), and had no effect on LNCaP cell growth in the absence or presence of hormone or IR (Figure 5J). Collectively, these data indicate that ZBTB10 overexpression is not sufficient to rescue enzalutamide radiosensitization or phenocopy the positive growth effect of androgen under IR conditions, possibly due to ZBTB10 not regulating DDR gene expression.

**Figure 5.**
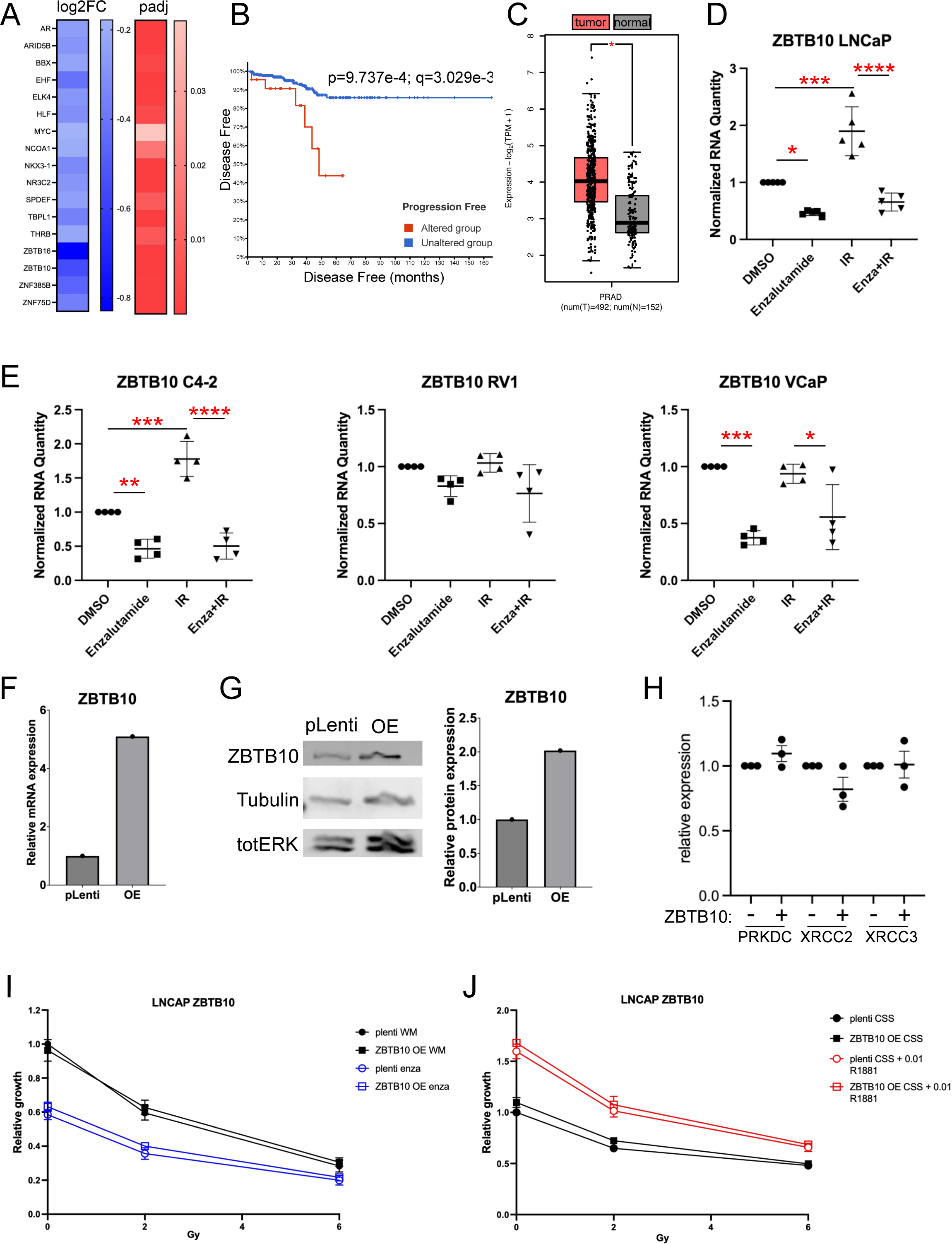
Effect of ZBTB10 overexpression on IR cell survival. (A) Log-transformed fold change and adjusted p-values of TFs repressed by enzalutamide. (B) Disease-free survival plot from the PCa TCGA showing a worse prognosis associated with alterations with ZBTB10 (q=0.041). (C) GEPIA2 analysis of the PCa TCGA and GTEx data of ZBTB10; *p<0.05. (D) LNCaP (E) C4-2, Rv1, VCaP normalized mRNA expression of ZBTB10 at 24 hours in response to 6Gy radiation, 10µM enzalutamide, or combination thereof. * <0.05, ** <0.005, *** <0.0005, **** <0.0001, using ANOVA followed by Sidak’s multiple comparisons test. (F, G) Validation of ZBTB10 overexpression showing mRNA (F) and protein (G) overexpression of LNCaP ZBTB10 stable mass population and vector control. (H) mRNA expression of in PRKDC, XRCC2, and XRCC3 in ZBTB10 overexpressing cells relative to control. (I, J) Relative growth over seven days in LNCaP ZBTB10 overexpression cells were irradiated with 0, 2, 4, or 6Gy and treated with vehicle control or 10µM enzalutamide (I) or 0.05nM R1881 (J).

## Discussion

There is compelling clinical evidence for the benefit of ADT in combination with RT (44–48). These clinical observations led to studies testing for interplay between the AR and the DDR (5,6). From these studies, the model that emerged was one of cooperation between the AR and DDR machinery; IR-induced DNA damage increases AR activation and transcription of DDR genes, and the AR is regulated by components of the DDR (5–10,12–16). However, specifics of this molecular model were inconsistent across the biological systems used in these studies, suggesting that aspects of this model of AR regulation of DDR genes were incomplete and in need of further examination. Furthermore, most of the studies leading to the model of AR regulating the DDR examined changes in transcription following treatment with hormone, anti-androgens, or IR after one to four days. Therefore, the transcriptional changes measured could be downstream and secondary from primary AR targets. Also, for the studies examining changes in transcription following hormone treatment, cells were often grown in charcoal-stripped serum containing media for one to three days; subsequent stimulation of these cells with hormone transitions cells from a quiescent state to an active growth state and thus, the transcriptional changes measured days later also encompasses growth associated transcript changes in addition to primary AR targets. We therefore sought to identify the immediate transcriptional changes induced by IR in an AR-dependent manner.

We found, counter to what is accepted in the field, that the AR is not a primary regulator of DDR genes. Using AR-positive androgen-responsive LNCaP PCa cells growing asynchronously in medium with whole serum, we measured transcriptional changes in gene expression using PRO-seq to quantify changes in nascent RNA transcription at early time points in response to IR, enzalutamide, and the combination of the two. Antagonizing the AR with enzalutamide significantly decreased expression of canonical AR target genes but had no effect on DDR gene sets. This observation is consistent with previously published GRO-seq data of hormone-stimulated LNCaP and VCaP cells, which did not identify any gene sets associated with the DDR (28,29). Examining individual DDR genes defined by previous studies as AR-regulated (7,8,10), we find only 6 of 75 genes significantly changed in response to enzalutamide. IR both increased and decreased canonical AR target gene transcription, and IR-responsive genes are not significantly closer to AR binding sites than IR-unchanged genes are, collectively indicating that the AR is not responsible for the primary IR-responsive transcriptional changes. When we explicitly test for genes activated by IR and repressed by enzalutamide, the pattern predicted for genes activated by AR in response to DNA damage, we find only 16 genes. None of these 16 genes are part of the Hallmark DNA Repair gene set. Furthermore, these genes do not have a consistent response to IR and enzalutamide across different PCa cell lines. Additionally, we did not observe changes in ELOVL7 protein levels in response to IR and enzalutamide treatment despite a robust, conserved, and persistent change in *ELOVL7* mRNA levels under the same experimental conditions in multiple PCa cell lines. Taken together, our data indicate that AR is not part of a primary IR-induced DNA damage transcriptional response in PCa cells, and AR is not a primary regulator of genes in the DDR under normal growth conditions. Therefore, the clinical benefit of combining ADT with RT is likely not due to AR directly regulating transcription of DDR genes in response to IR-induced DNA damage.

The DDR protein – AR interactions are relatively underexplored and could generate a delayed AR transcriptional response in reaction to IR. We measured nascent transcription following 90 minutes of enzalutamide treatment (30 minutes enzalutamide alone -/+ 60 minutes IR). Interactions of DDR proteins with the AR could lead to primary AR transcriptional changes with delayed kinetics. In response to IR, gH2AX and 53BP1 foci peaked at 2hrs, and when we examined DNA damage using the comet assay, we found that DNA damage was mostly repaired in PCa cells by 2 hours (18). These kinetics suggest that we should have observed a primary AR transcriptional response to IR under the conditions we used here if AR was regulating expression of genes important for DNA repair through DDR protein – AR interactions. If the AR is a critical regulator of the DDR through steady state regulation of genes in the DDR, then one possibility is that this regulation represents a secondary effect mediated through one or more of the 17 TFs directly regulated by the AR. One of these TFs, ZBTB10, is amplified in 8% of PCa in the TCGA dataset, and in 26% in the East Coast SU2C metastatic CRPC (mCRPC) dataset (49,50). However, overexpression of ZBTB10 alone was not sufficient to replace the protective effect from IR afforded by androgen treatment or induce expression of three putative AR-regulated DDR genes, PRKDC, XRCC2, or XRCC3. Interestingly, we observed MYC downregulation in response to enzalutamide, suggesting MYC transcript levels are directly regulated by AR in LNCaP cells. This is in contrast to what was observed in VCaP cells (28) at 2hrs DHT stimulation, suggesting AR regulation of MYC in PCa may be heterogeneous. VCaP cells express the TMPRSS2-ERG fusion (51) and TMPRSS2-ERG can regulate MYC expression levels (52), raising the possibility that MYC may be coordinately regulated by AR and ERG. Overexpression of MYC, in the context of normal AR expression levels, clearly downregulates AR transcriptional programs through coregulator redistribution and Pol II promoter-proximal pausing (53–55), suggesting a negative feedback loop between AR and MYC. What is still unknown is how AR and MYC transcriptional programs interact at steady state when the relative endogenous expression levels of both MYC and AR are considered (55).

In this study we also examined Pol II localization to determine the impact of IR and AR inhibition on Pol II initiation and promoter-proximal pausing. Interestingly, enzalutamide treatment led to an increase in promoter-proximally paused Pol II. In earlier studies, we found that CDK9, which associates with cyclin T to form pTEFb (56), phosphorylates AR on Ser81 (31). This phosphorylation is required for optimal AR transcriptional activity and association with chromatin, is the highest stochiometric phosphorylation on the AR, and continues to increase over time with hormone treatment (30,31,57,58). Our observation of enzalutamide treatment increasing Pol II promoter-proximal pausing lends further support for the model that AR regulates transcription through a feedforward loop involving mobilization of pTEFb, increasing AR pS81 phosphorylation, and co-regulator recruitment that actively sustains the transcription of AR-regulated genes (59).

The data presented here imply that the clinical benefit of ADT in combination with RT is not due to direct AR transcriptional regulation of the DDR. If AR antagonism limits DNA repair through reducing expression of DDR genes, as predicted by the integrated model proposed from previous publications (7,8,10,11), then neoadjuvant ADT should be more effective than adjuvant ADT when combined with RT. However, the opposite seems true; adjuvant ADT is more effective than neoadjuvant ADT in combination with RT (48,60). Thus, we propose that the clinical benefit of ADT and RT is not due to the primary AR regulation of DDR gene transcription. The field needs to consider alternative mechanisms for the clinical benefit of combining ADT with RT, including non-tumor cell-autonomous mechanisms. AR clearly has important roles in many cell types (61). Effects on the tumor microenvironment (TME) of combining ADT with RT could lead to clinical benefit. Furthermore, non-tumor cell-autonomous mechanisms accounting for the clinical benefit of combining ADT with RT could explain the variability among PCa tumor cell lines to respond to IR. Recent studies have shown that androgen signaling suppresses T-cell immunity against cancer (62,63) raising the possibility that ADT counteracts this immunosuppression, which may be critical to any abscopal effect induced by RT (64,65). Other possible mechanisms could account for the ADT plus RT benefit, including the straightforward possibility that both treatments independently reduce tumor cell fitness.

## Methods

### Cell culture

LNCaP (RRID:CVCL_0395) and C4-2 (RRID:CVCL_4782) cells (gifts from Dr. L. W. K. Chung) and C4-2 CAMKK2 knockout cells generated previously (39) were grown in DMEM:F12 (Invitrogen) with 5% non-heat inactivated serum (Gemini) and 1% Insulin-Transferrin-Selenium-Ethanolamine (ITS) (ThermoFisher). CWR22Rv1 (22Rv1) (RRID:CVCL_1045; gift from Dr. Steven Balk) and VCaP (RRID:CVCL_2235; gift from Dr. Kerry Burnstein) were grown in DMEM (Invitrogen) with 5% heat-inactivated serum. Commercial DNA fingerprinting kits (ATCC) verified cell lines and all cell lines were tested for mycoplasm.

### Cell treatment for gene expression analysis

To determine which sites are actively transcribed in PCa cells upon IR treatment and to see if these changes depend on the AR, we performed two independent PRO-seq experiments with cells treated as follows. For all experiments, LNCaP cells were plated into 150 mm tissue-culture dishes at 4.2 x 10^6^ cells, grown two days in the whole medium under standard growth conditions (37°C, 5% CO_2_). Cells were then treated with 1) mock or 2) 6 Gy IR followed by 60 min incubation for Exp 1 (Figure 1A). For Exp 2 (Figure 2A), cells were treated with 1) DMSO (control) vehicle for 90 min; 2) DMSO vehicle for 30 min plus 6 Gy IR followed by 60 min incubation; 3) 10 µM enzalutamide for 90 min; and 4) 10 µM enzalutamide for 30 min followed by 6 Gy IR and incubated for 60 min. Cells were permeabilized as previously described (35). These early time points were selected to identify the primary transcriptional effects and are consistent with the kinetics of steroid uptake, AR antagonism, and DNA damage/repair.

### PRO-seq

PRO-seq library preparation and analysis were performed as previously described (66–68). In brief, cells were permeabilized according to Mahat et al. (2016) (35). The permeabilized cells were counted, and 50 µL aliquots containing approximately 6-8 x 10^6^ cells were snap-frozen in liquid nitrogen and stored at −80°C. PRO-seq libraries were prepared according to Sathyan et al. (2019) (66), which included a random eight-base unique molecular identifier (UMI) at the 5′ end of the adapter ligated to the 3′ end of the nascent RNA. No size selection was performed to avoid bias against short nascent RNAs.

### qPCR

RNA isolation and quantitative real-time PCR (qPCR) were performed as previously described (69,70). RNA concentrations were determined using a NanoDrop 2000 UV-Vis Spectrophotometer (Thermo Scientific). cDNA was synthesized using Sensifast cDNA synthesis kit (Bioline, BIO-65054). Quantitative real-time PCR (qPCR) was performed using iTaq Universal SYBR Green Supermix (Biorad, #1725124). Primer sequences: CAMKK2 F: CGTCTCCATCACGGGTATGC, CAMKK2 R: CACCATAGGAGCCCTTTCCA; CDC42EP3 F: GCATTGGTGATGTTCGGCTC, CDC42EP3 R: GCCCTTCATAGCTAGTGCCA; ELOVL7 F: GTTTGGGAACATTCCATGCCC, ELOVL7 R: ATGATGCACGCAAAGACTGG; PMEPA1 F: CTGGTTCAGAGAAGGCCGAG, PMEPA1 R: GAGGACAGGGAATGAACCCG; PVT1 F: TGGAATGTAAGACCCCGACTCT; PVT1 R: GATGGCTGTATGTGCCAAGGT, TLE1 F; GATCTGGGGATCACGCCTTG, TLE1 R: ACCACTGCTCTACAAAGGACG; WNT7B F: CAAAGTCTGCCAGCAACAGG; WNT7B R: CAGCTTTTATTGGGCCGAGC. The relative standard curve method was used to determine transcriptional fold changes as we have done previously (71–74). RNA starting quantities (SQ) were determined using a standard curve. The SQ mean was then normalized to the reference gene PSMB6.

### Western blot analysis

Western blotting performed as previously described (69,70). Antibodies: ELOVL7 (Sigma AV49908; used at 1:1000), ZBTB10 (Bethyl Laboratories A303-257A; used at 1:5000), ERK (Cell Signaling 137F5; used at 1:1000), Ran (BD Biosciences #610340; used at 1:2000), and Tubulin (Cell Signaling #2144; used at 1:1000).

### Generation of ZBTB10 overexpressing stable cells

Reverse transduction was done by adding plenti-HA virus or pLenti-ZBTB10-HA virus diluted to an MOI of 25 in DMEM-F12 media containing 5% non-heat inactivated FBS and 1x ITS to a fibronectin (1 μg/ml) coated 6 well plate and then adding LNCaP cells to the well. Two days after the reverse transduction, media was changed and 1 μg/ml puromycin selection was added. Cells were fed with selection media every two days until reaching 80% confluency. The cells were then expanded with continued selection until freeze-downs could be made for future propagation and experimentation.

### Growth assay

Assay was performed as previously described (18). Briefly, shRNAs targeting ELOVL7 (sh#1 sense: ccggtGTTACTTCTCCAAATTTAAtacctgacccataTTAAATTTGGAGAAGTAACTTTTTg; sh#2 sense: ccggtCCTGGTGGTTTGGAGTCAAATCTCGAGATTTGACTCCAAACCACCAGGTTTTTg), or WNT7B (sh#1 sense: ccggtGCATGAACCTGCATAACAAtacctgacccataTTGTTATGCAGGTTCATGCTTTTTg; sh#2 sense: ccggtGTGGCAGTGCAACTGCAAATTctcgagAATTTGCAGTTGCACTGCCACTTTTTg), or Vector control virus was added to fibronectin-coated (1 μg/ml) 96-well plates, or LNCaP ZBTB10-HA stable cells or LNCaP vector cells were plated in triplicate at 3000 cells/well in 96-well plates, one plate per irradiation condition, either in phenol-red free DMEM-F12 media containing 5% non-heat inactivated FBS and 1x ITS for enzalutamide condition or phenol-red free DMEM-F12 media containing 5% charcoal-stripped FBS and 1x ITS for the R1881 condition. Immediately following plating, 0.05 nM R1881 was added to cells designated to receive hormone stimulation. 24 hours after plating and 30 minutes prior to irradiation, 10 μM enzalutamide was added to cells designated to receive the inhibitor. The cells were treated with 0, 2, 6 Gy of radiation. CyQuant Direct reagent was added on day 6 according to the manufacturer’s protocol (ThermoFisher). Quantification was performed using a BioTek Synergy 2 plate reader.

### Data Availability Statement

The raw PRO-seq sequencing files can be accessed from the NCBI Gene Expression Omnibus (GEO; https://www.ncbi.nlm.nih.gov/geo/query/acc.cgi?acc=GSE235902) under accession number GSE235902. All analysis details and code are available at https://gioeli-lab.github.io/AR_IR_PRO-seq_analysis/Vignette.html.

