## Supplemental Files for "The androgen receptor does not directly regulate the transcription of DNA damage response genes"

**Supplemental Figure 1. MA plots.** (A) MA plot corresponding to volcano plot in Figure 1D. (B) MA plot of PRO-seq signal in dREG defined putative regulatory elements in response to IR. (C) MA plots corresponding to volcano plots in Figure 2F. (D) MA plots corresponding to volcano plots in Figure 2G. (E) MA plots corresponding to volcano plots in Figure 2H. (F) MA plots corresponding to volcano plots in Figure 2I.

**Supplemental Figure 2. Expression and biology of differentially regulated gene subsets.** (A-C) Normalized count expression data for genes (A) activated by radiation and enzalutamide, (B) repressed by radiation and enzalutamide, and (C) activated by enzalutamide and repressed by radiation. (D) Over-representation analysis of the genes responsive to the treatments at an FDR of 0.05. (E-F) Kaplan-Meier curves for progression-free survival from the PCa TCGA patients with and without alterations in the 16 genes positively regulated by the AR and IR from Figure 3A, and (H) the subset of the top eight genes activated by IR and repressed by enzalutamide. Alterations here are inclusive of mutations, structural variants, copy number alterations, and mRNA expression differences with a z-score threshold of 2.0 relative to all samples. Comparable significance was found when mRNA expression changes were excluded.

**Supplemental Figure 3. Regulation of candidate AR-regulated DDR genes in PCa cell lines**. Normalized mRNA expression of CAMKK2, CDC42EP3, ELOVL7, PMEPA1, PVT1, TLE1, and WNT7B at 24 hours in response to 6Gy radiation, 10μM enzalutamide, or combination thereof in C4-2, 22Rv1, and VCaP cells. (A) CAMKK2, (B) CDC42EP3, (C) ELOVL7, (D) PMRPA1, (E) PVT1, (F) TLE1, (G) WNT7B. * <0.05, ** <0.005, *** <0.0005, **** <0.0001 indicating statistically significant effect of enzalutamide on control or IR, using ANOVA followed by Sidak's multiple comparisons test.

**Supplemental Figure 4. Effect of ELOVL7 and WNT7B knockdown and CAMKK2 knockout on cell growth.** Relative growth over seven days in (A, D) LNCaP ELOVL7 knockdown, (B, E), LNCaP WNT7B knockdown, and (C, F) C4-2 CAMKK2 knockout cells grown in whole media without and with (A-C) 10µM enzalutamide or in CSS without and with (D-F) 0.05nM R1881. * <0.05, ** <0.005, *** <0.0005, **** <0.0001, using ANOVA followed by Sidak's multiple comparisons test. (G) Confirmation of CAMKK2 knockout and ELOVL7 and WNT7B knockdown.

**Supplemental Figure 5. Effect of ELOVL7 and WNT7B knockdown and CAMKK2 knockout on cell survival after IR.** (A-F) Relative cell survival over seven days in (A, B) LNCaP ELOVL7 knockdown, (C, D) LNCaP WNT7B knockdown, and (E, F) C4-2 CAMKK2 knockout cells were irradiated with 0, 2, 4, or 6Gy and treated with vehicle control or 0.05nM R1881 (A, C, E) or 10µM enzalutamide (B, D, F).

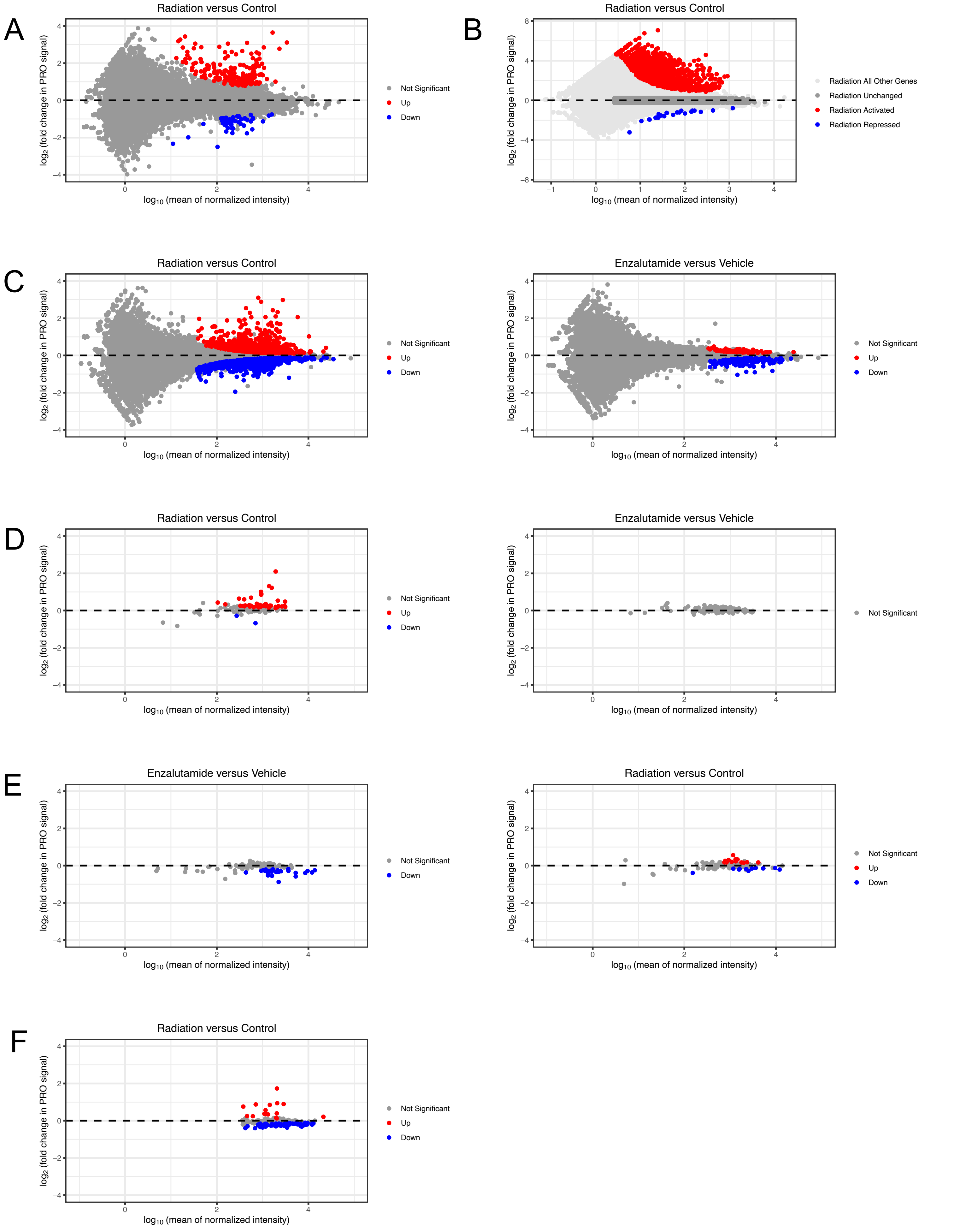


Supplemental Figure 1

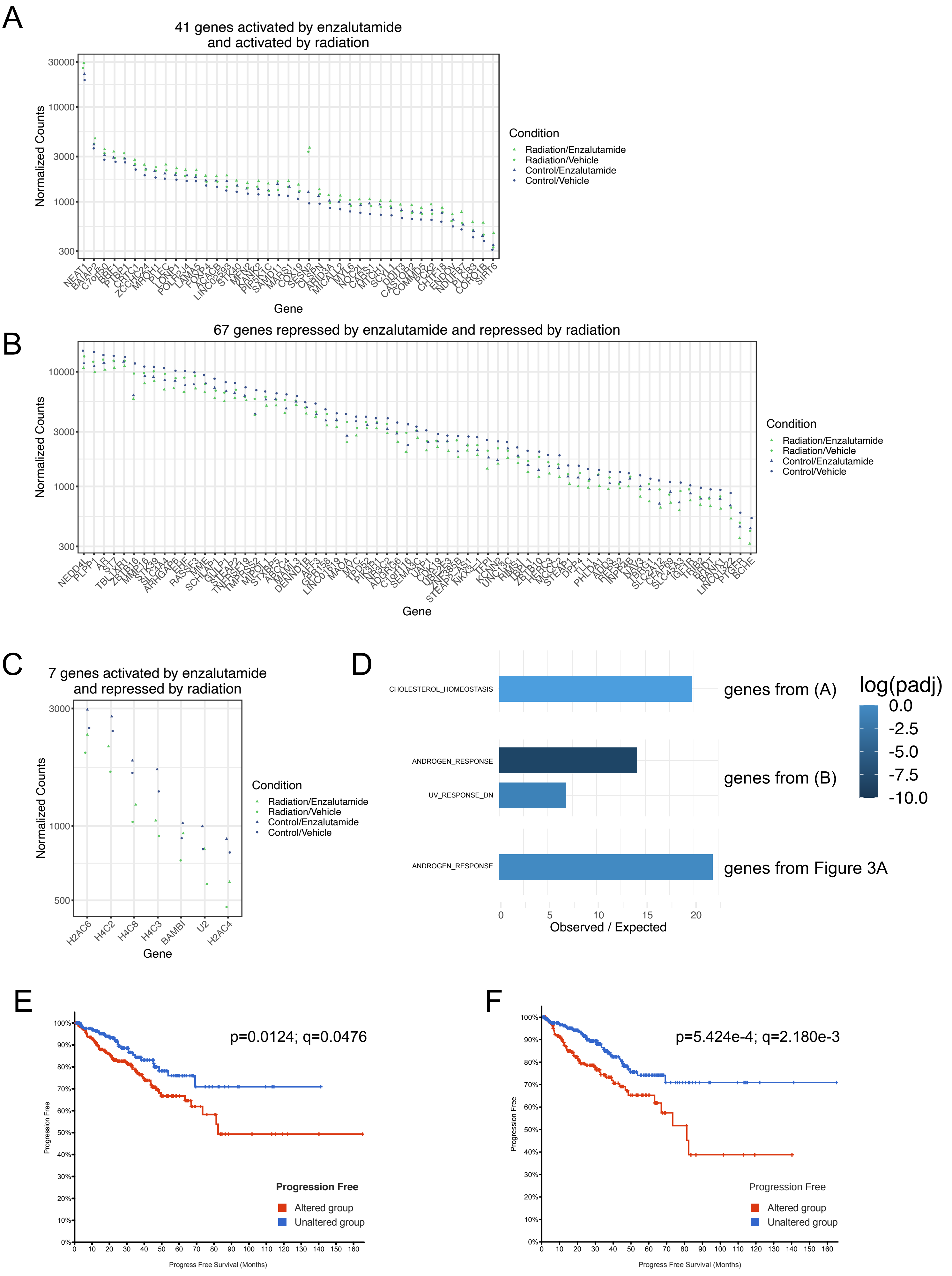


Supplemental Figure 2

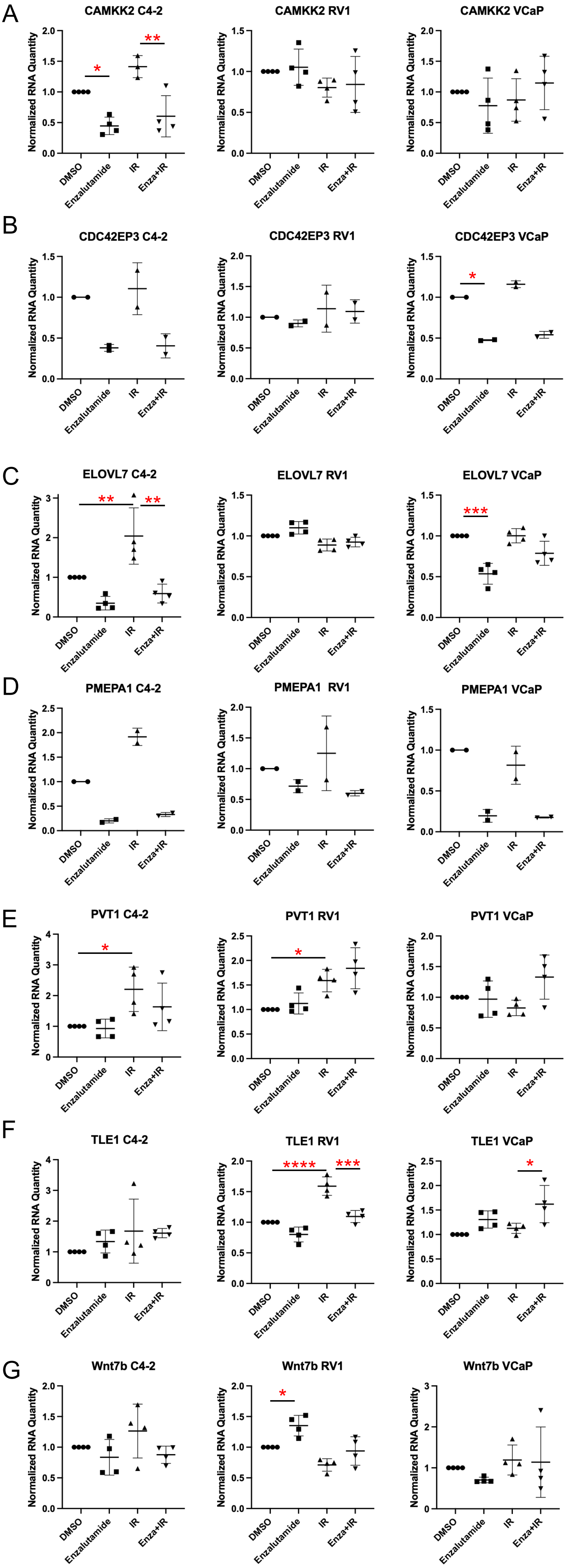


Supplemental Figure 3

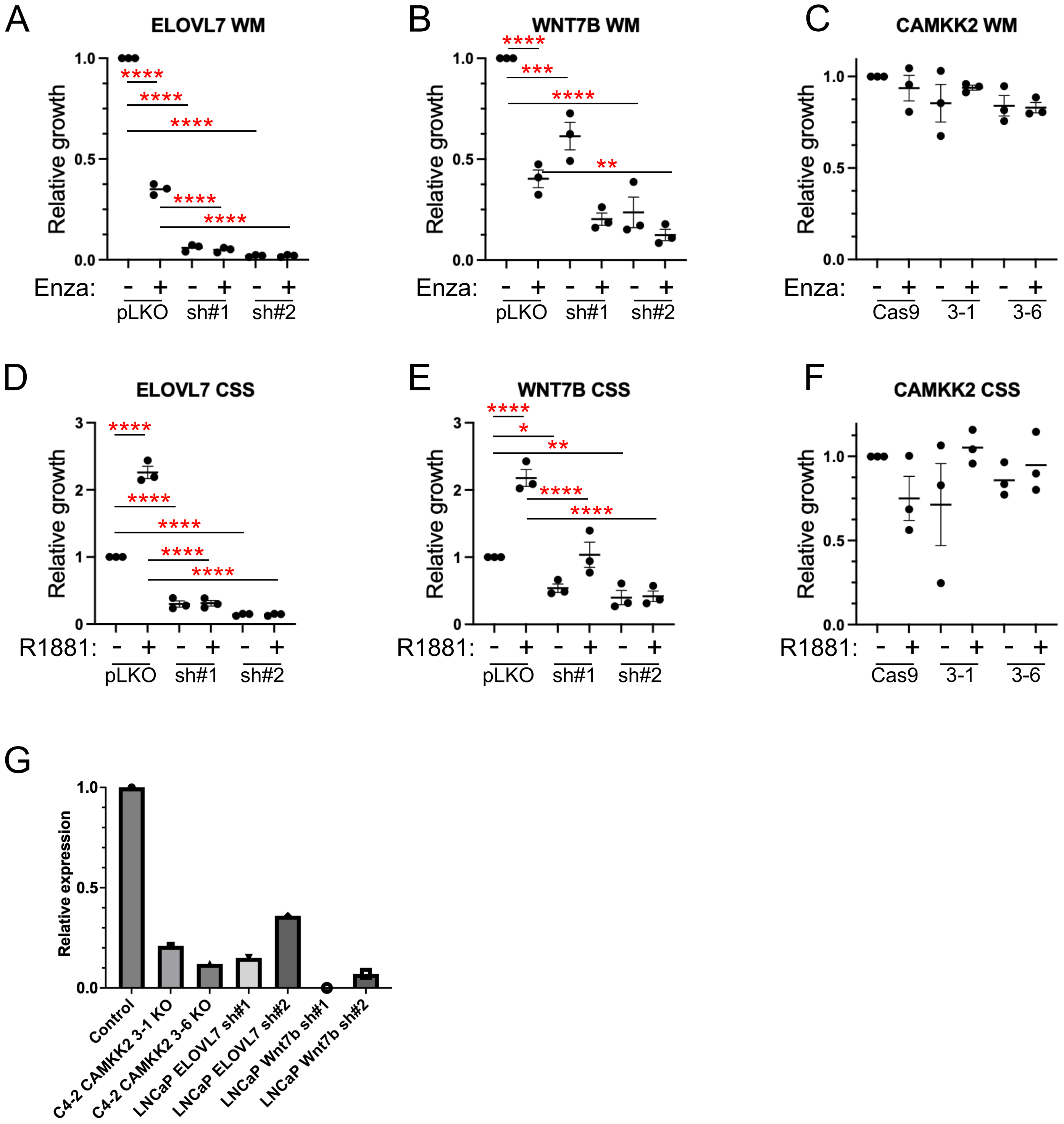


Supplemental Figure 4

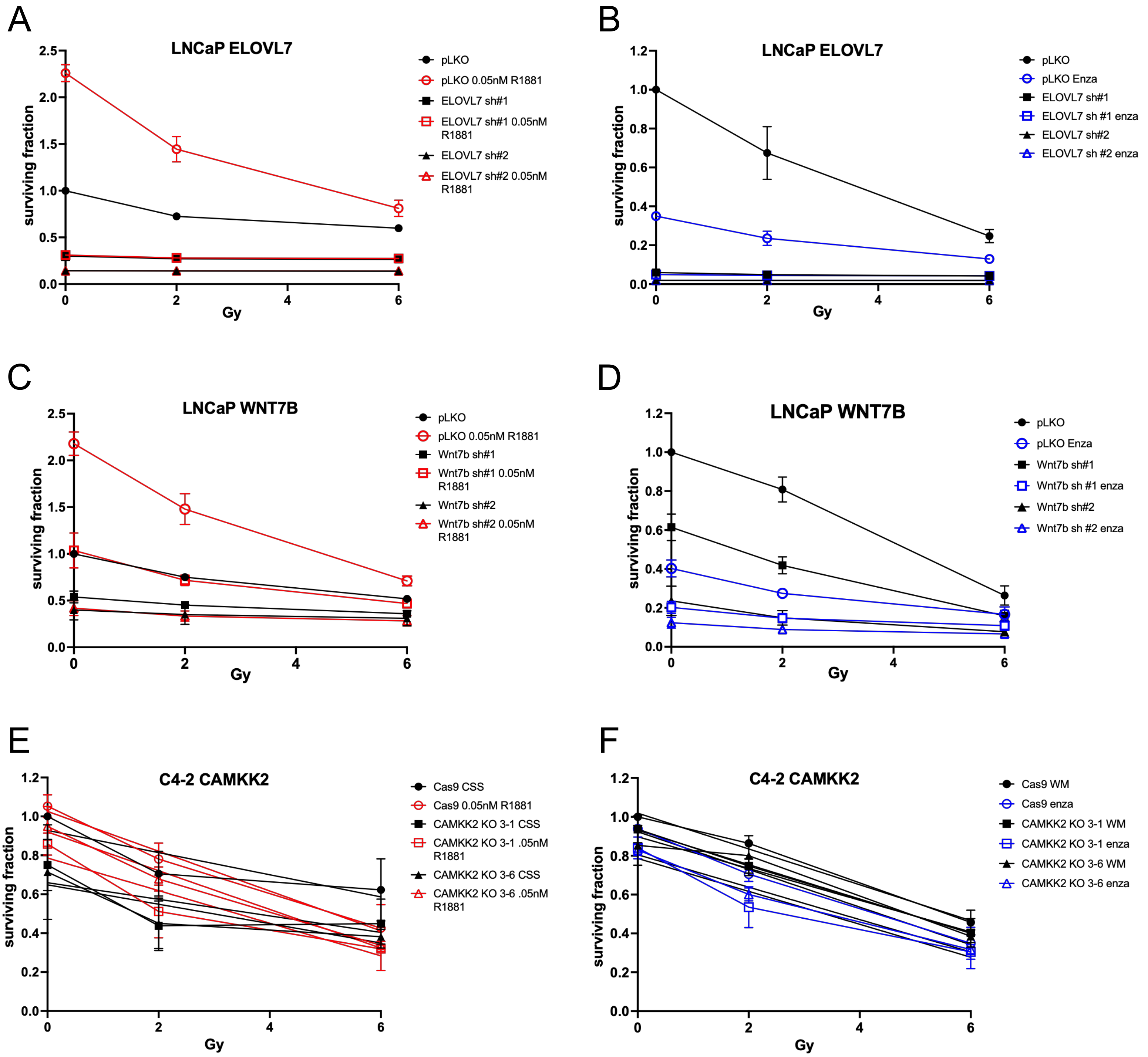


Supplemental Figure 5
